# Spinal Fusion Properties of Mechanically-Reinforced, Osteomodulatory Chitosan Hydrogels

**DOI:** 10.1101/2022.05.26.493540

**Authors:** Blake T. Darkow, Joseph P. Herbert, Mark J. Messler, Abigail Grisolano, August J. Hemmerla, Austin D. Kimes, Julien Lanza, Yisheng Sun, Julia R. Crim, Derek Stensby, Caixia Wan, Don K. Moore, Bret D. Ulery

## Abstract

Lower back pain is a considerable medical problem that will impact 80% of the U.S. population at some point in their life. For the most severe cases, surgical repair is necessary and is associated with costs upwards of $10.2 billion annually in the United States. To alleviate back pain, spine fusions are a common treatment in which two or more vertebrae are biologically fused together often through the use of a graft material. Unfortunately, iliac crest bone autograft, the current gold standard graft material, can yield insufficient fusion and is associated with considerable donor site morbidity and pain as well as limited supply. Therefore, new materials need to be developed in order to better coordinate healing and new bone growth in the affected area to reduce unnecessary patient burden. In order to address this issue, the incorporation of allograft and one of two types of cellulose (*i.e*., ^0^CNCs and CNFs) into a dual-crosslinked chitosan hydrogel loaded with bioactive calcium phosphate was investigated. Hydrogels were then tested for both their material and biological properties. Specifically, hydrogel swelling ratio, mass loss, ion release profile, compressive strength, *in vitro* biocompatibility and osteoinduction as well as *in vivo* biocompatibility, and effectiveness in a spine fusion model were determined. Cellulose and allograft incorporation significantly improved hydrogel compressive strength and biocompatibility and CNFs were found to be a significantly more biocompatible form of cellulose than ^0^CNCs. Additionally, through the controlled delivery of osteoinductive simple signaling molecules (*i.e*., calcium and phosphate ions), DCF-loaded CNF/Chitosan hydrogels were able to induce osteoblast-like activity in murine mesenchymal stem cells. When evaluated *in vivo*, these hydrogels were found to be non-toxic though the subacute phase (14 days). A 6-week rabbit spine fusion found these materials to achieve near complete fusion when assessed radiographically. This research provides considerable support for the utility of our novel material for spine fusion procedures as well as other future bone applications.

## Introduction

Lower back pain is a considerable medical problem impacting 80% of the United States population at some point in their life.^1,2^ It is the second most common reason for doctor’s visits in the United States and greatest cause of workplace absence in the U.K.^3^ While only a minority of the most severe cases require surgery, they account for 29.3% of the total expenditures associated with lower back pain.^4^ Reports on the total healthcare spending of these procedures vary widely, from $784 million to $10.2 billion annually in the United States.^4,5^ Regardless of which figure is accurate, the burden of lower back pain on society is immense.

Conventional treatment for lower back pain involves an escalation of invasiveness starting with conservative options such as physical therapy before progressing to less-invasive surgeries such as disc repair and replacement. If these approaches do not address a patient’s symptoms, then spinal fusion may be necessary.^3^ Spinal fusion procedures aim to alleviate back pain brought on by preexisting conditions through mechanically and biologically fixing two or more adjacent vertebrae through the use of instrumentation and/or bone graft materials. The total volume of fusion procedures only continues to rise (from 164,527 in 2004 to 281,575 in 2015) as the U.S. population ages.^5^ Despite the significant increase in cases, the rate of fusion achieved clinically varies widely among different bone graft materials from as low as 40% to as high as nearly 100%.^5,6^ Additionally, the current gold standard for graft material (*i.e*., iliac crest bone autograft) is associated with considerable donor site morbidity and pain as well as limited supply.^7^ The significant drawbacks associated with the currently available treatment options as well as increases in the number of procedures performed annually highlights the need for new bone graft substitutes to be developed.

Recent efforts by the biomaterials community have been focused on utilizing tissue engineered scaffolds to mimic the physical, chemical, and biological constructs that exist in natural tissues in order to coordinate healing. Our research groups have focused on the utilization of simple signaling molecules to influence the differentiation of select cell populations to regenerate tissues of interest. For spinal fusion applications, we have developed an osteoinductive biomaterial comprised of a dual-crosslinked, cellulose-supported chitosan hydrogel loaded with bioactive calcium phosphate.^8-10^ This hydrogel is designed to release calcium and phosphate ions within a previously defined therapeutic window in order to induce osteoinduction in mesenchymal stem cells. This desirable bioactivity was previously achieved by the dissociation of a phosphate crosslinker^8^ and later by controlled release from dibasic calcium phosphate.^9^ While this has laid the groundwork for the use of this biomaterial for spinal fusion procedures, there remains significant work to be conducted to continue the development of this hydrogel before it can be motivated to the clinic.

The primary aim of this research is to expand the knowledge surrounding materials for bone tissue engineering applications. Specifically, improvements to the osteoinductive, cellulose-reinforced chitosan hydrogel developed in this lab will be made with focus on its utilization in lumbar fusion procedures to treat lower back pain. Despite significant progress in the development of this material, our previously published work falls short of reaching the *in vitro* mechanical and biocompatible benchmarks necessary to be studied *in vivo* in appropriate animal models. For example, the compressive strength of the current formulation is lower than that of natural bone and the hydrogels have been found to suppress the proliferation of mesenchymal stem cells which are crucial for bone regeneration. Through this research, a further engineered solution has been generated that possesses more desirable mechanical and biocompatibility properties which was evaluated both *in vitro* and *in vivo*.

## Materials and Methods

### Formation of Chitosan Hydrogels

#### Chitosan/Cellulose Solution

To create a chitosan solution, low molecular weight chitosan (m.w. 50,000 - 190,000 Da, Sigma Aldrich) was dissolved in 0.5% acetic acid supplemented deionized, distilled water (ddH_2_O) at 1.4% weight/volume and magnetically stirred for 3 days at room temperature followed by gravity filtration through cotton (Fisher Scientific). To mechanically reinforce the chitosan hydrogels, cellulose was chosen as a material. Neutral cellulose nanocrystals (*i.e*., ^0^CNCs) were processed from cellulose microcrystals using a previously described method.^8,9^ Cellulose nanofibrils (i.e., CNFs) were purchased from the Process Development Center at the University of Maine. CNFs were produced by mechanical grinding wood pulp until fibers approximately 20 – 50 nm in width and several hundred microns in length were produced.^11^ ^0^CNCs or CNFs were added to the chitosan solution at 0.07% weight/volume and dispersed using 10 minutes of sonication via a probe-tip sonicator (Branson Sonifer 450) at 20 W. For osteoinductive ion releasing hydrogels, 6% and/or 10% weight/volume dibasic calcium phosphate (DCP_6_, DCP_10_, Jost Chemical Co.) were added to the chitosan/cellulose solution. The DCP was then dispersed using 1 minute of sonication via the probe-tip sonicator at 20 W.

#### Crosslinker Solution

A dual crosslinking solution was created by dissolving 42 mg/mL of sodium bicarbonate (*i.e*., ionic crosslinker) and 28 mg/mL of genipin (*i.e*., covalent crosslinker) in ddH_2_O followed by 1 minute of sonication via the probe tip sonicator at 20 W. The crosslinking molar ratio was 5:1.25:1 of carbonate : genipin : deacetylated chitosan site (∼ 85%). This ratio was chosen based on the advantageous mechanical and bioactive properties previously published with this ratio.^9^

#### Hydrogel Formation

Hydrogels were formed by adding the appropriate amount of chitosan/cellulose solution to crosslinker solution at a ratio of 5:1 volume/volume followed by 1 second of high-speed mixing on a vortex mixer. The gelation vials were then placed in a 37 °C water bath while they were monitored every 30 seconds for their gelation time. Hydrogels were determined to have gelated upon the material remaining adhered to the bottom of the gelation vials when inverted.^12^ After gelation was verified, the hydrogels were allowed to set in a 37 °C incubator for 24 hours before further testing. For allograft-loaded hydrogels, 20% weight/volume crushed cancellous allograft (AG, 0.1 – 4 mm, MTF Biologics) was placed in the gelation vial before the chitosan/cellulose solution and then the crosslinker solution were then added.

### Physical Characterization of Hydrogel

#### Swelling Ratio

Hydrogel swelling was determined by first submerging preformed hydrogels in phosphate buffered saline (PBS, Gibco) at 37 °C for 24 hours. After 24 hours, the samples were removed from PBS and excess water was removed from the surface by Kim wipe blotting before they were weighed. The hydrogels were then frozen at - 80 °C and lyophilized under vacuum (0.1 mm Hg) at - 50 °C for 72 hours to ensure all solvent was removed. After lyophilization, the hydrogels were weighed again and swelling ratio was determined using **Equation 1**:

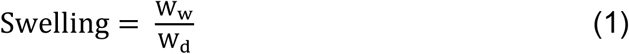

where W_w_ is the wet hydrogel weight before lyophilization and W_d_ is the dry hydrogel weight after lyophilization.

#### 3.2.2 Mass Loss

Hydrogel mass loss was measured by submerging the samples in phosphate buffered saline (PBS) at 37 °C. At specific intervals, the samples were removed from solution and excess water was removed from the surface by Kim wipe blotting before they were weighed. After weighing, the hydrogels were gently submerged in the PBS again. Mass loss was calculated using **Equation 2**:

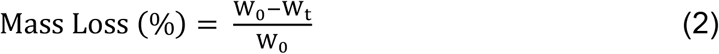

where W_0_ is the initial weight and W_t_ is the weight at a specific time point. Ion Release

Calcium (Ca^2+^) and phosphate (PO_4_^3-^) ions released from DCP and DCP-loaded hydrogels were measured in ddH_2_O at 37°C over 14 days. Preformed hydrogels were immersed in ddH_2_O in 24 well plates and DCP powder was placed into 24-well semi-permeable inserts (Greiner Bio-One). At specific timepoints, ddH_2_O for each sample was exchanged and assayed for ion content. Ca^2+^ release was evaluated using the Calcium (CPC) LiquiColor™ Test (Stanbio Laboratory). Release samples (1 - 10 μL) were combined with 95 μL of base and color reagent and mixed. The resultant solution was then read at 550nm using a BioTek Cytation 5 fluorospectrometer. Ca^2+^ ion concentrations were determined by comparing the sample readings to a standard curve (0 - 10 mM). PO_4_^3-^ release was measured using a Phosphate Colorimetric Assay (Sigma). Release samples (1 - 100 μL) were combined with 30 μL of assay reagent and diluted with ddH_2_O until a final volume of 200 μL was obtained. Solution absorbance was measured at 650 nm and compared to a standard curve (0 - 5 μM) in order to determine the PO_4_^3-^ concentration in the samples.

#### Compressive Strength

Preformed cylindrical hydrogels (height ∼ 5 mm and diameter ∼ 12 mm) were carefully removed from their gelation vials. Their compressive strength was determined via compression testing using an Instron universal tester. Hydrogels were compressed at a rate of 1 mm/min until 80% strain was achieved at which point the strength was calculated by dividing the maximum force by the compressed hydrogel surface area.

### Assessment of *In Vitro* Cytotoxicity and Bioactivity

#### Cell Culture

Murine mesenchymal stem cells (mMSCs) were purchased from Cyagen and stored in the vapor phase of a liquid nitrogen dewar. mMSCs were seeded into T-75 cell culture flasks (CytoOne) and cultured at 37°C in a humidified incubator with 5% CO_2_ in complete growth media consisting of Dulbecco’s modified Eagle’s medium (DMEM, Gibco) supplemented with 10% fetal bovine serum (FBS, Sigma) and 1% penicillin-streptomycin (Pen-Strep, Gibco). Media was changed every 48 hours until the cells reached ∼ 80% confluency at which time they were dissociated using a 0.05% trypsin-EDTA solution (Gibco). Delaminated mMSCs were then counted using a hemocytometer and passed into new T-75 flasks at a splitting ratio of 1:5. Surplus mMSCs were cryopreserved in complete growth media supplemented with 10% DMSO. After the fifth passage, cells were used for *in vitro* studies.

#### Acute Cytotoxicity

Tissue culture treated 24-well plates (CytoOne) were seeded with 100,000 cells/well and exposed to complete growth media as a negative control. Preformed chitosan hydrogels with no cellulose, ^0^CNCs, or CNFs (*i.e*. C:G:C 5:1.25:1, ^0^CNC/C:G:C 5:1.25:1, and CNF/C:G:C 5:1.25:1) were washed twice with PBS and then soaked in 1 mL of complete growth media for 24 hours to create extract media. The extract media was then transferred to the seeded 24 well plates in which the cells were cultured for 24 hours at 37°C in a humidified incubator at 5% CO_2_ after which cell quantification and viability assays were conducted on the samples.

#### Cell Proliferation Assay

Cell proliferation was determined using a Quant-iT™ PicoGreen™ dsDNA Assay (Invitrogen). At each timepoint, cells were washed with PBS and lysed using 1% Triton X-100 (Sigma-Aldrich) followed by three freeze-thaw cycles and sonication via a probe tip sonicator (Branson Sonifer 250) at 10 W to fully lyse the cells. Lysates were diluted with TE buffer (200 mM Tris-HCL, 20 mM EDTA, pH 7.5) and then mixed with PicoGreen reagent according to the manufacturer’s recommended protocol. The fluorescence of each sample was read using a BioTek Cytation 5 fluorospectrometer (*ex*. 480 nm, *em*. 520 nm), and cell number was determined using a mMSC standard curve (0 - 250,000 cells/mL).

#### Cell Viability Assay

Cell viability was evaluated using an alamarBlue™ cell viability reagent (Invitrogen) assay. At each timepoint, all experimental groups were removed from the wells and the cells were gently washed with PBS. The cells were then cultured in complete growth media supplemented with 10% alamarBlue™ reagent for 1 hour at 37 °C in a humidified incubator at 5% CO_2_. After incubation, the fluorescence of the alamarBlue™ media for each sample was measured (*ex*. 560 nm, *em*. 590 nm). Cell viability was reported as a ratio of the emission in the experimental groups compared to the group exposed to the complete growth media negative control.

#### Anti-Abrasion Platform for Inductivity Assessment

A novel anti-abrasion sample platform (**Figure 1**) was developed to remove the impact mechanical stress and diffusion limitations would have if hydrogels were placed directly on top of plated cells. These platforms were printed using an SLA 3D printer (Form 2, Formlabs) and washed using 91% isopropyl alcohol to remove any residual unbound resin. Samples were washed twice with PBS and soaked in complete growth media for 24 hours before being place into wells to support the hydrogel samples during *in vitro* studies. Each group, including non-hydrogel controls, contained a platform in order to reduce the influence the platform’s footprint had on cells.

**Figure 1.**
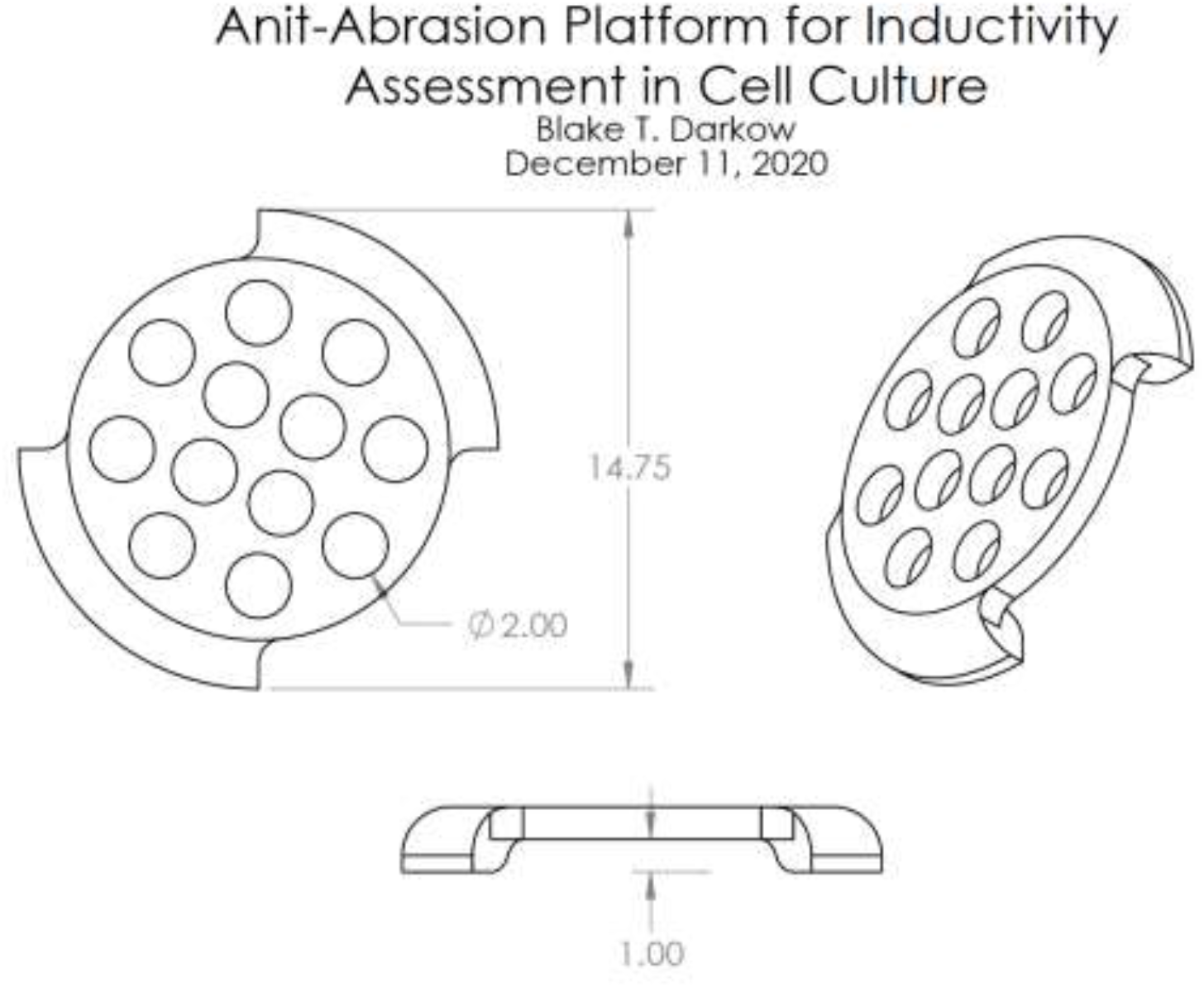
Anti-Abrasion Platform for Inductivity Assessment Cell Culture. The raised platform is perforated to allow mass transfer through and from the biomaterial in question while alleviating the adherent cells from mechanical stress due to contact. Dimensions are in millimeters.

#### *In Vitro* Bioactivity

Tissue culture treated 24-well plates (CytoOne) were seeded with 50,000 cells/well and exposed to complete growth media as a negative control. Osteogenic media was created by supplementing complete growth media with 2 mM L-glutamine, 50 μM ascorbic acid, 0.1 μM dexamethasone, and 10 mM β-glycerophosphate. CNF/C:G:C 5:1.25:1 hydrogels alone or with DCP_10_ and/or AG were washed twice with PBS and once with complete growth media. The hydrogels were then gently placed on an in-lab fabricated 3-D printed anti-abrasion platform (Formlabs) in 24-well plates with seeded cells and co-cultured at 37°C in a humidified incubator at 5% CO_2_ for which the growth media was changed every two days throughout the study. Cell quantity, viability, alkaline phosphatase activity, and mineralization were evaluated at days 3, 7, 14, and 21.

#### Alkaline Phosphatase Activity Assay

Cell alkaline phosphatase (i.e., ALP) activity was measured at each time point using an Alkaline Phosphatase Activity kit (BioVision). In brief, 80 μL of cell lysate (harvested as described for the proliferation assay) was combined with 50 μL of 5 mM *p*-nitrophenyl phosphate (pNPP) in assay buffer. The reaction was incubated for 1 hour at room temperature and then stopped by adding 20 μL of stop solution (NaOH). The absorbance of the resulting solution was read at 405 nm using a BioTek Cytation 5 fluorospectrometer. Absorbance values were converted to enzyme activity by comparing them to a standard curve (0 - 20 μM) of dephosphorylated pNPP (*i.e*., pNP). ALP activity was reported as pNP content normalized to cell count.

#### Cell-based Mineralization Assay

Cell-based mineral deposition was measured using an Alizarin Red assay. At each time point, cells were gently washed with ddH_2_O and fixed in 70% ethanol for 24 hours. The ethanol was removed and the cells were covered with 1 mL of 40 mM Alizarin Red solution (Sigma-Aldrich) for 10 minutes. The samples were then gently but thoroughly washed with ddH_2_O to remove all unbound stain. Absorbed Alizarin Red stain was then desorbed using a 10% cetylpyridinium chloride (CPC, Sigma-Aldrich) solution which was harvested and its absorbance measured at 550 nm via a BioTek Cytation 5 fluorospectrometer. Absorbance readings were then converted to Alizarin Red concentration using a standard curve (0 - 0.274 mg/mL). The same procedure was performed on acellular 24-well plates that had undergone the same experimental conditions and these values subtracted from the cellularized samples to calculate cell-based mineralization. All values reported were normalized by cell count.

### Assessment of *In Vivo* Biocompatibility and Bioactivity

#### Subacute Biocompatibility

All animal research was conducted with approval of the Animal Care and Use Committee at the University of Missouri - Columbia. Subacute biocompatibility was assessed by a 14-day intramuscular implantation model utilizing C57BL/6J mice (Jackson Laboratories). CNF / C:G:C hydrogels were prepared with and without 10% DCP (DCP_10_) and with or without crushed cancellous chip allograft (AG) (N = 4). Additionally, a sterilization method was evaluated and compared to hydrogels formed as previously described. For the additional sterilization step, the CNF/ C:G:C solution was first autoclaved at 121 °C for 30 minutes and crosslinker solution sterile filtered (0.22 μm) before combining to generate the final product. A single injection of 50 μL saline served as a negative control for the procedure. Mice were anesthetized with Ketamine / Xylazine (100 mg/kg / 10 mg/kg) and appropriate analgesia was provided every 12 hours as needed via Buprenorphine (0.05 - 0.1 mg/kg). Following confirmation of anesthetic depth via the absence of a toe pinch reflex, the hind limb was shaved and aseptically prepped. A 5 - 10mm incision was then made followed by blunt dissection to create a pocket of uniform depth within the underlying biceps femoris muscle. The implant material (50 μL) was placed within the freshly made pocket and the incision was closed with absorbable sutures whereas the skin was closed with surgical adhesive (VetOne). Mice were given adequate supportive care and monitored daily for 7 days after surgery for clinical symptoms of implant rejection. After 14 days, the animals were euthanized and their implantation sites were checked for signs of infection. The tissue encompassing the implant was then removed and fixed in 10% formalin for histological staining.

#### Histology Scoring

Tissue samples were fixed in 10% formalin for 48 hours followed by paraffin embedding, microtome sectioning, and hematoxylin and eosin staining by IDEXX Laboratories. Slides were then individually evaluated across all groups and scored by a pathologist. Examination criteria for histological evaluation of tissue sections was determined using the scoring scale detailed in ISO 10993 – 6 “Biological evaluation of biomedical devices part 6 – test for local effects after implantation”. The specifics of this scoring scale are presented in **Table 1**.

**Table 1.** Examination criteria for histological evaluation of tissue sections. Scoring scale is according to ISO 10993 – 6 “*Biological evaluation of biomedical devices part 6 – test for local effects after implantation*”. This table is reprinted with permission from previously published work.^13^

| Cell Type / Response | Score |  |  |  |  |
| --- | --- | --- | --- | --- | --- |
|  | 0 | 1 | 2 | 3 | 4 |
| Polymorphonuclear cells | 0 | Rare, 1 - 5/phf* | 5 - 10/phf | Heavy infiltrate | Packed |
| Lymphocytes | 0 |  |  |  |  |
| Plasma cells | 0 |  |  |  |  |
| Macrophages | 0 |  |  |  |  |
| Giant cells | 0 | Rare, 1 - 2/phf | 3 - 5/phf |  | Sheets |
| Necrosis | 0 | Minimal | Mild | Moderate | Severe |
| Neovascularization | 0 | Minimal capillary proliferation, focal, 1 to 3 buds | Groups of 4 to 7 capillaries with supporting fibroblastic structures | Broad band of capillaries with supporting fibroblastic structures | Extensive band of capillaries with supporting fibroblastic structures |
| Fibrosis | 0 | Narrow band | Moderately thick band | Thick band | Extensive band |
| Fatty infiltrate | 0 | Minimal amount of fat associated with fibrosis | Several layers of fat and fibrosis | Elongated and broad acculation of fat cells around the implant site | Extensive fat completely surrounding the implant |
| Traumatic necrosis | 0 | Minimal | Mild | Moderate | Severe |
| Foreign debris | 0 |  |  |  |  |

\*phf = per high-powered (400x) field

#### Spine Fusion Procedure

To evaluate the potential for chitosan hydrogels to address back pain, a New Zealand White Rabbit spinal fusion model was utilized.^14,15^ During this procedure, CNF / DCP_10_ / C:G:C and AG / CNF / DCP_10_ / C:G:C hydrogels were compared to the implantation of AG alone (N = 6). All hydrogels were sterilely formed as previously described before implantation. After formation, the hydrogels were lyophilized (Labconco) for 48 hours. Rabbits (3 - 4 kg) were premedicated with 0.025 mg/kg of Dexmedetomidine and 10 mg/kg of Ketamine delivered intramuscularly. Buprenorphine SR (0.2 mg/kg) was given subcutaneously to provide prolonged analgesia. Anesthesia during the procedure and/or 2D radiography was provided with 2% - 4% Isoflurane inhalant. The lumbar region on the back was shaved and rabbits were placed on the operation table in the prone position. The surgical site was prepped using 3 alternating washes with chlorohexidine and isopropyl alcohol wipes. A lidocaine injection was made into the operation area for local anesthesia. To begin the procedure, two fascial incisions parallel to the midline between the multifundus and longissimus muscles will be made near the L5 and L6 vertebrae. Blunt dissection was then used to access the transverse processes at the fusion level where 0.5 mL of gentamicin (1 mg/mL) was sprayed into the prospective fusion site on each side. The inside faces of the adjacent transverse processes were decorticated using a surgical drill (Stryker™ Core Sumex™). The graft material was then placed between the transverse processes. The wound and subcutaneous tissue were sutured using 3-0 absorbable suture (Securos Surgical Securosorb™). The superficial skin layer was closed using 4-0 absorbable suture (Ethicon™ Monocryl) with buried knots. Pulse, temperature, and respirations were recorded at least every 15 minutes until the animal was ambulatory. The rabbits were then closely monitored to ensure surgical site healing, infection prevention, appropriate analgesia, and return to normal activity for up to 10 days post-operation. After 6 weeks, the rabbits were euthanized via intravenous injection of Euthasol (0.22 mL/kg) and the lumbar spine was harvested and fixed in 10% Formalin.

#### Rabbit Radiographs and Scoring

Lumbar spine 2D radiographs were taken prior and immediately after as well as 3 and 6 weeks after the spinal fusion operation was performed. Radiographs were taken in the anterior – posterior, posterior – anterior, and lateral views. Radiographs were taken on a Innovet Classic Model E739x at 52 mA / 25 kVp and digitally scanned via an AGFA CR 30-X plate reader. To evaluate the 2D Radiographs, they were independently analyzed following the conclusion of the trials by human clinical evaluators blinded to the experimental groups. Each timepoint was scored according to the scale displayed in **Table 2**. The score for each animal at each timepoint is presented as the average score between all evaluators where standard deviation was calculated as the standard deviation among the average scores.

**Table 2.** Scoring system for evaluation of spine fusion via 2D Radiograph.

|  |  |
| --- | --- |
| 0 | Bone graft resorption, no new bone formation |
| 1 | New bone formation between but not spanning the transverse processes |
| 2 | Bone spanning the transverse processes without fusion |
| 3 | Partial fusion |
| 4 | Complete fusion |

### Statistical Analysis

JMP software was used to make comparisons between experimental groups with Tukey’s HSD test specifically employed to determine pairwise statistical differences (p < 0.05). Groups that possess different letters have statistically significant differences in mean, whereas those that possess the same letter have means that are statistically insignificant in their differences.

## Results and Discussion

The osteoinductive hydrogel developed in our lab utilizes components that are individually understood to be biocompatible molecules and materials. Even so, the lack of proliferation in MSCs exposed to our hydrogels remains an area requiring further investigation.^8,9^ A potential source of MSC suppression is the surface modification of CNCs with cetyltrimethylammonium bromide (CTAB) to produce neutral CNCs (^0^CNCs). To establish a more biocompatible solution to SSM delivery via a chitosan-based hydrogel, this research explored alternatives to the novel ^0^CNCs and their impact on the material properties and biocompatibility of the hydrogel.

### Impact of Agent Incorporation on Chitosan Hydrogel Material Properties

Surgical biomaterials that are formed or cured *in situ* require valuable time that must be used judiciously. Therefore, a 20-minute window for a formulation to turn from liquid to gel has been defined.^16^ Gelation times for different chitosan hydrogels (*i.e*., Carbonate:Genipin:Chitosan 5:1.25:1 - C:G:C) varied slightly based on their DCP content and cellulose type (**Figure 2**). The addition of DCP_x_ (*i.e*., DCP_6_ or DCP_10_) to the hydrogels resulted in a faster gelation time for DCP_x_ / C:G:C and CNF / DCP_x_ / C:G:C hydrogels. The effect of DCP on expediting gelation time may be due to the excess of potential crosslinking agents available because of reversable ionic bonds being formed through the phosphate content of DCP. This finding is consistent with our previously published data.^9^ However, DCP incorporation did not yield faster gelation times with either ^0^CNC / DCP_x_ / C:G:C hydrogels suggesting that the ^0^CNCs limit or inhibit DCP mediated-crosslinking. This effect may be due to the hydrophobicity of ^0^CNCs reducing the overall generation of additional network crosslinks by DCP. All formulations gelled within the desired 20 minute window making these formulations viable options for time-sensitive *in situ* gelation in a surgical setting.^16^

**Figure 2.**
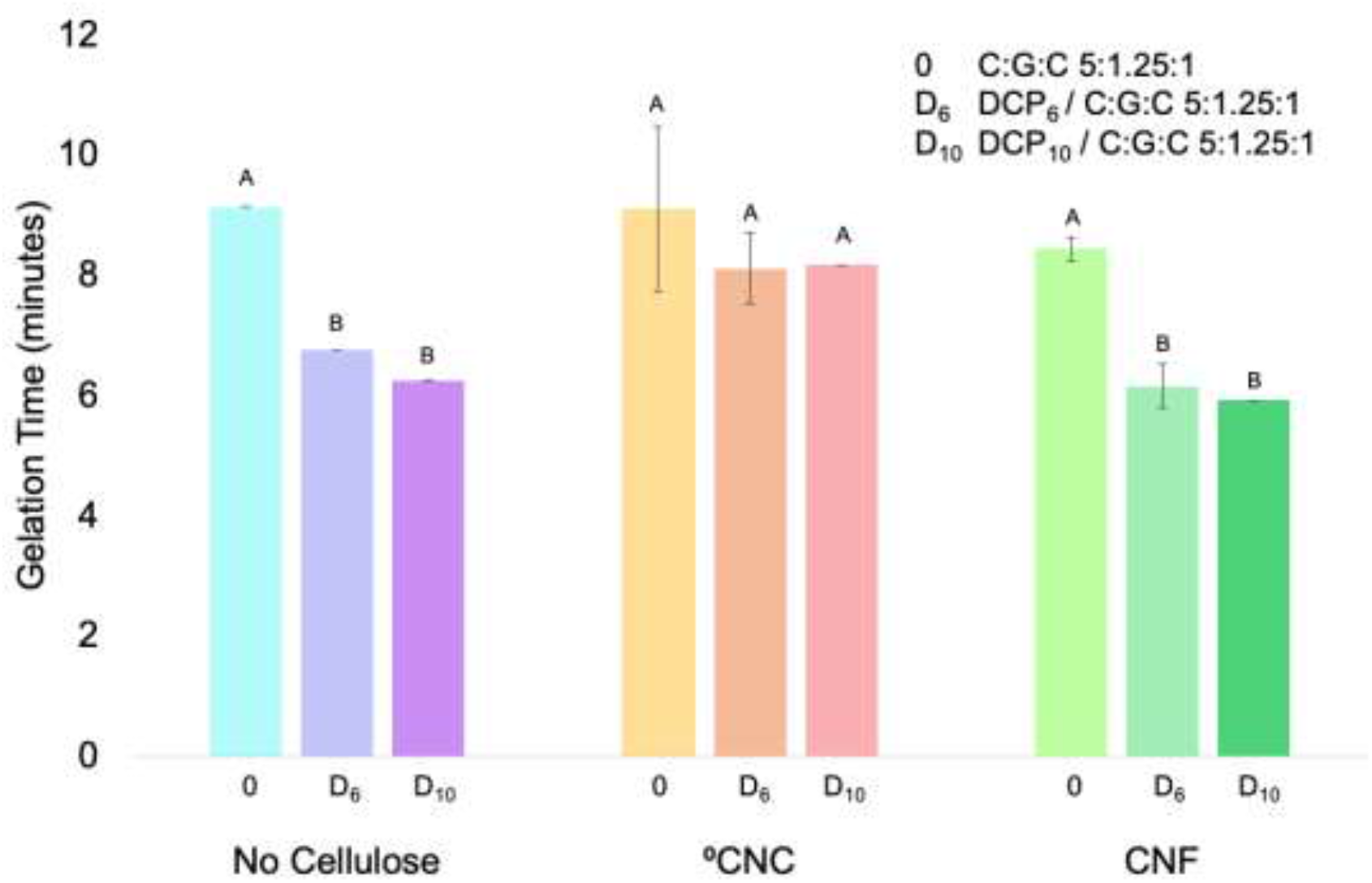
Gelation times for C:G:C 5:1.25:1 hydrogels formed with various cellulose types and DCP concentrations. Hydrogels were prepared with combinations of no cellulose, ^0^CNCs, or CNFs, without DCP (0), with 6% DCP (D_6_), or with 10% DCP (D_10_). The inversion method at 37 °C was used to determine *in situ* gelation time. Values are reported as the average ± standard deviation (N = 4). Statistical groupings are based on a Tukey’s HSD test between all groups. Groups that possess different letters have statistically significant differences (p < 0.05) in their means whereas those that possess the same letter are statistically similar.

While gelation time is one important factor for biomaterial utility, its swelling ratio can provide valuable insight into porosity and hydrophilicity. To evaluate the swelling capacity of the hydrogels, the gels were immersed in PBS for 24 hours at 37 °C and then lyophilized. Hydrogel wet weight (pre-lyophilization) and dry weight (post-lyophilization) were then used to determine the swelling ratio (**Table 3**). The swelling ratio for all hydrogels decreased significantly with the incorporation of DCP. This is likely due to the increase in physical density and crosslinking generated by the inclusion of CaP. The two types of cellulose investigated had different impacts on the swelling ratio compared to the C:G:C hydrogel alone. ^0^CNC incorporation decreased the swelling ratio across all gel formulations though only statistically significantly for the hydrogels lacking DCP (*i.e*., C:G:C). A possible explanation for this observation is that the component which gives ^0^CNCs their neutral surface charge, CTAB, possesses a long alkyl chain (16 hydrocarbons in length) which creates a more hydrophobic environment due to inter- and intra-molecular hydrophobic interactions. Conversely, CNFs increased the swelling ratio for the DCP incorporated hydrogels though only statistically significantly for the DCPr_6_ group. As the CNFs used in this study are mechanically grinded from wood pulp with no surface modifications,^11^ they are relatively hydrophilic in nature,^17^ making it unsurprising that CNF incorporation would increase the swellability of the chitosan hydrogel.

**Table 3.** Swelling ratios for C:G:C 5:1.25:1 hydrogels with various cellulose types and DCP concentrations. This was determined as the wet weight divided by the dry weight after immersion in PBS at 37 °C for 24 hours. Values are reported as the average ± standard deviation (N = 4). Statistical groupings are based on a Tukey’s HSD test between all groups. Groups that possess different letters have statistically significant differences (p < 0.05) in their means whereas those that possess the same letter are statistically similar.

|  | C:G:C | DCP<sub>6</sub> / C:G:C | DCP<sub>10</sub> / C:G:C |
| --- | --- | --- | --- |
| No Cellulose | 14.55 ± 0.94 (A) | 5.76 ± 0.19 (D) | 4.73 ± 0.15 (D,E) |
| $^0\text{CNC}$ | 10.95 ± 0.28 (B) | 4.83 ± 0.09 (D,E) | 3.94 ± 0.09 (E) |
| CNF | 15.48 ± 0.79 (A) | 7.73 ± 0.26 (C) | 5.74 ± 0.44 (D) |

Another important characteristic of an implantable biomaterial is its ability to retain its components while guiding the body through the healing process. With this in mind, mass loss for the hydrogels was investigated over a one-week period with the day 1, 3, and 7 mass retention ratios presented in **Figure 3**. The largest drop in mass was seen within the first day in which the hydrogels lost 9 - 21% of their mass and the two days after that in which the hydrogels were relieved of another 10 - 20% of their mass. However, beyond day 3, the hydrogels stabilized, like due to the more loosely bound contents having completely dissociated by this time point. When cellulose alone was incorporated into the hydrogels, no differences were observed in mass loss regardless of the formulation. Interestingly, when both cellulose and DCP were incorporated, ^0^CNCs facilitated statistically significantly greater mass retention at days 3 for both DCP concentrations (*i.e*., 6 wt. % and 10 wt. %), but only 6 wt. % DCP at day 7. These data are similar to results previously published with ^0^CNC / DCP_x_ / C:G:C hydrogels.^9^ When DCP begins to dissociate, some PO_4_^3-^ ions can remain entrapped in the hydrogel matrix due to their anionic nature facilitating additional ionic crosslinking molecules with open primary chitosan amines available in the network. However, for the hydrogels with no cellulose or with CNFs incorporated, this was not observed. One potential explanation for this result is that, based on the swelling ratio of the respective hydrogels, the no cellulose and CNF hydrogels swell to a much greater extent decreasing their solid mass content and increasing their porosity. This effect limits hydrogel component retention, likely preventing the additional crosslinking possible with DCP generated PO_4_^3-^ and increasing the availability for mass transfer out of the hydrogel as it dissociates.

**Figure 3.**
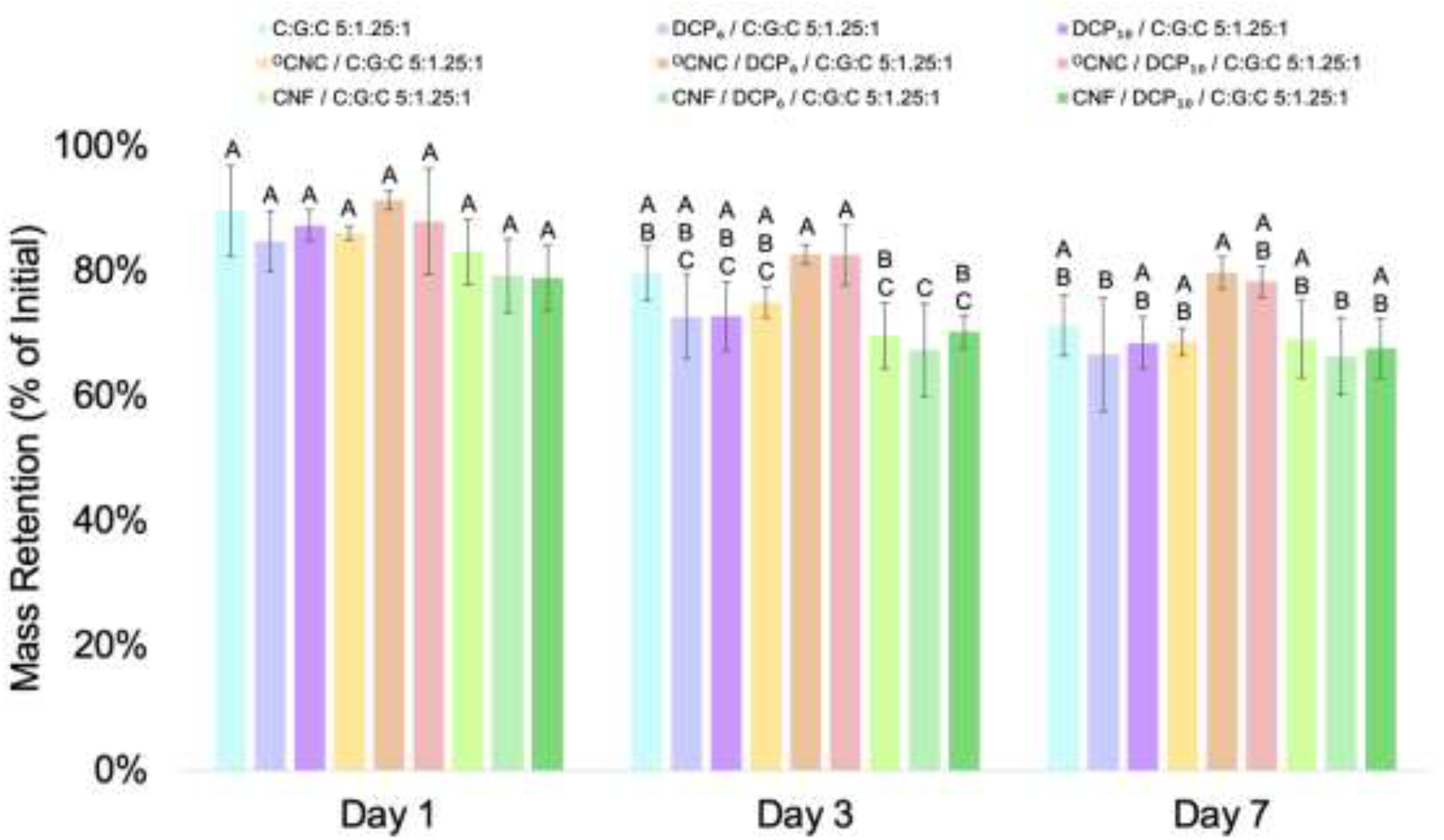
Mass retention for C:G:C 5:1.25:1 hydrogels with various cellulose types and DCP concentrations. This was determined after immersion in PBS at 37 °C for up to 7 days. Values are reported as the average ± standard deviation (N = 4). Statistical groupings are based on a Tukey’s HSD test between all groups. Groups that possess different letters have statistically significant differences (p < 0.05) in their means whereas those that possess the same letter are statistically similar.

In addition to mass loss by itself being an important factor when considering biomaterial choice, product dissociation also impacts its capacity to facilitate controlled SSM delivery. Release profiles of Ca^2+^ and PO_4_^3-^ from DCP and DCP-loaded chitosan hydrogels are presented in **Figure 4**. The overall ion release profile was found to be similar regardless of cellulose type and DCP content. Ca^2+^ release was relatively consistent throughout the 14 days of the experiment (**Figure 4A**). In contrast, hydrogels released the greatest amount of PO_4_^3-^ within the first 24 hours (**Figure 4B**). This initial burst of PO_4_^3-^ may have been a result of the hydrogel network having the greatest carbonate-based crosslinking density at the beginning of the study. As evident in the mass loss data, the hydrogel dissociates over time likely freeing up chitosan cationic amines allowing for the negatively charged PO_4_^3-^ to remain associated with the positively charged polymer network. This phenomenon also explains why the Ca^2+^ release was much greater than that of the PO_4_^3-^, even though they have equivalent molar ratios within DCP (*i.e*., chemical formula - CaHPO_4_). The ion concentrations generated by the dissociation of DCP more or less remains within the previously determined osteoinductive window (*i.e*., 1 - 16 mM for Ca^2+^ and 1 - 8 mM for PO_4_^3-^) where they are both non-cytotoxic and osteoinductive.^10^ In fact, the ion levels never reach the toxic concentrations of 32 mM Ca^2+^ / 16 mM PO_4_^3-^ nor the non-bioactive concentrations of 0.5 mM Ca^2+^ / 0.5 mM PO_4_^3-^.^10^

**Figure 4.**
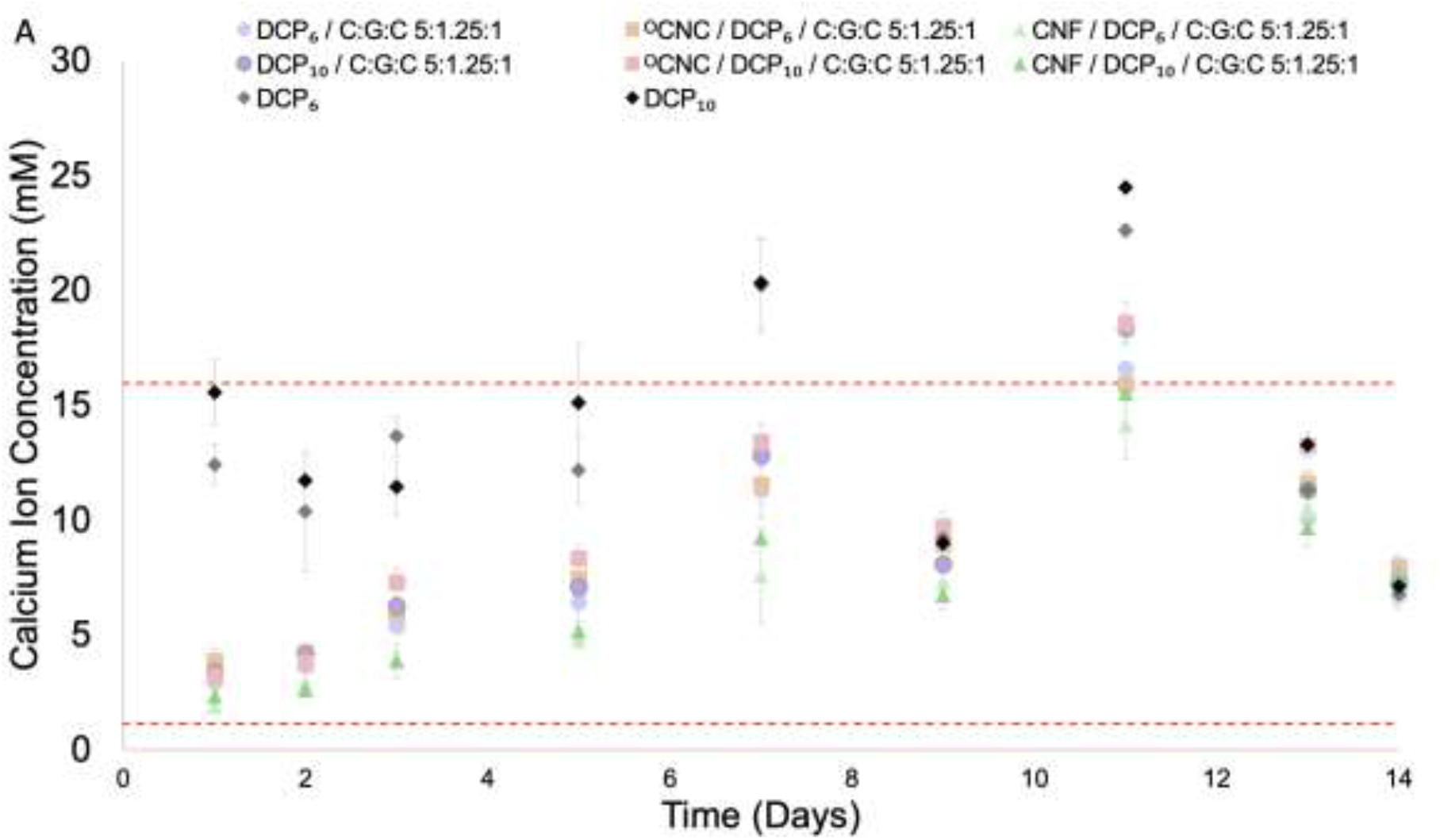
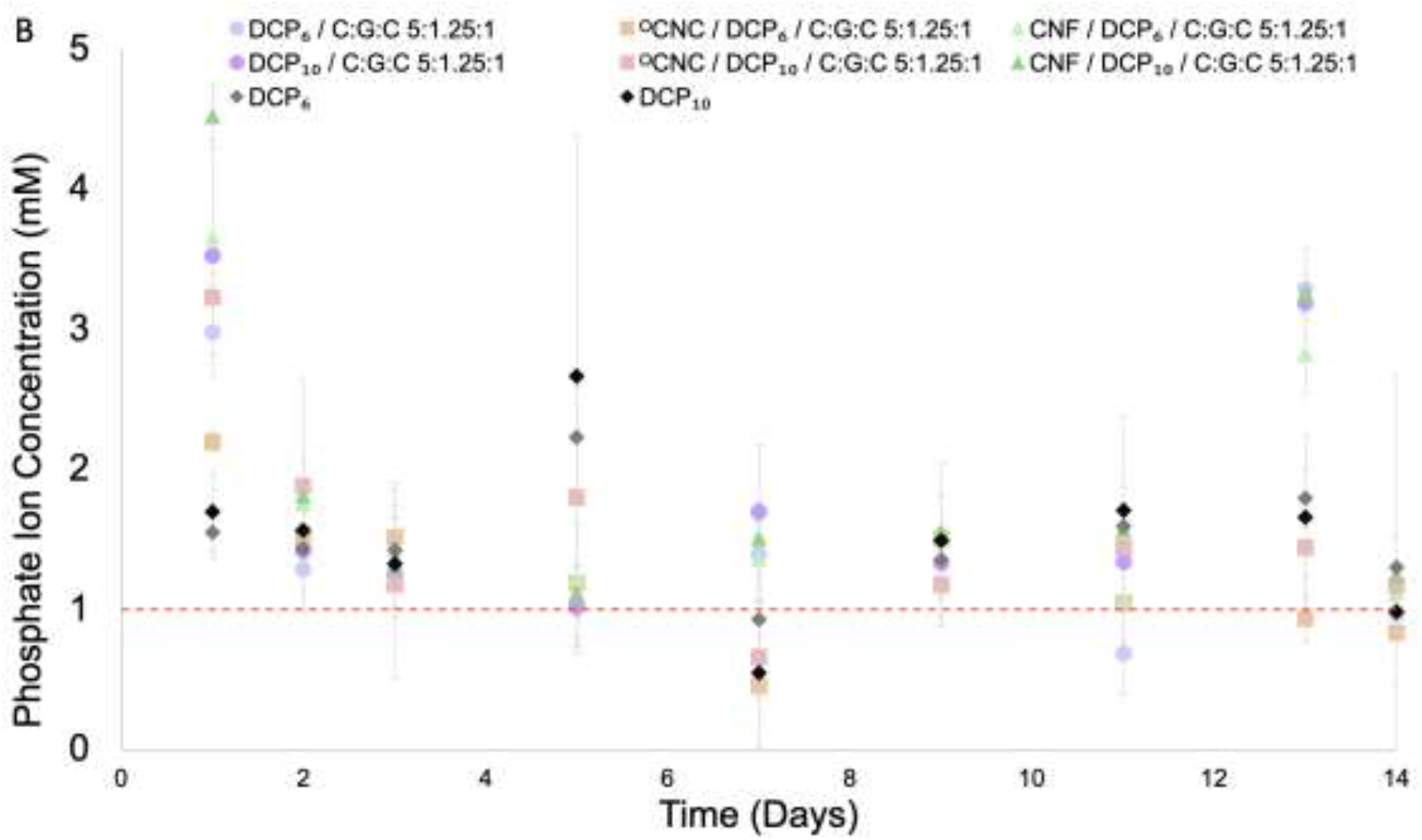
Calcium ion (Ca^2+^) and phosphate ion (PO_4_^3-^) release concentration from C:G:C 5:1.25:1 hydrogels formed with various cellulose types and DCP concentrations. Hydrogels were prepared with combinations of no cellulose, ^0^CNCs, or CNFs, without DCP (0), with 6% DCP (D_6_), or with 10% DCP (D_10_). **(A)** Ca^2+^ and **(B)** PO^4^_3-_ release from hydrogels immersed in ddH2O was measured at 37 °C for 14 days. Values are reported as the average ± standard deviation (N = 4). The Ca^2+^ concentration bounds and PO^4^_3-_ concentration lower bound of the osteoinductive therapeutic windows are shown with a red line.

While the ion release profile of the implanted material governs its osteoinductivity, the material must also possess the necessary compressive strength to be suitable for weight bearing bone regeneration. The compressive strength of these self-supported hydrogels are displayed in **Figure 5**. The addition of cellulose to the hydrogel without DCP incorporation did result in a slight increase in compressive strength, although this improvement was not statistically significant. Excitingly, the effect of DCP incorporation was highly impactful, especially within the ^0^CNC and CNF groups. The addition of ceramics like CaP to hydrogels has been widely reported to improve their mechanical strength.^9,18^ As previously discussed, phosphate from DCP can interact with the amine sites on chitosan yielding even greater crosslinking enhancing the mechanics of the material. Independently, ^0^CNCs and CNFs have also been specifically shown to improve the mechanical strength of hydrogels.^8,19^ Nanomaterials have been found to improve the mechanical strength of hydrogels by interlocking neighboring fibers with one another providing additional mechanical support.^19^ The synergistic effect observed with DCP and cellulose co-incorporation may be attributable to the overall increased mass density of the hydrogels as well as the cooperative, and not competitive, mechanically reinforcing nature of each component.

**Figure 5.**
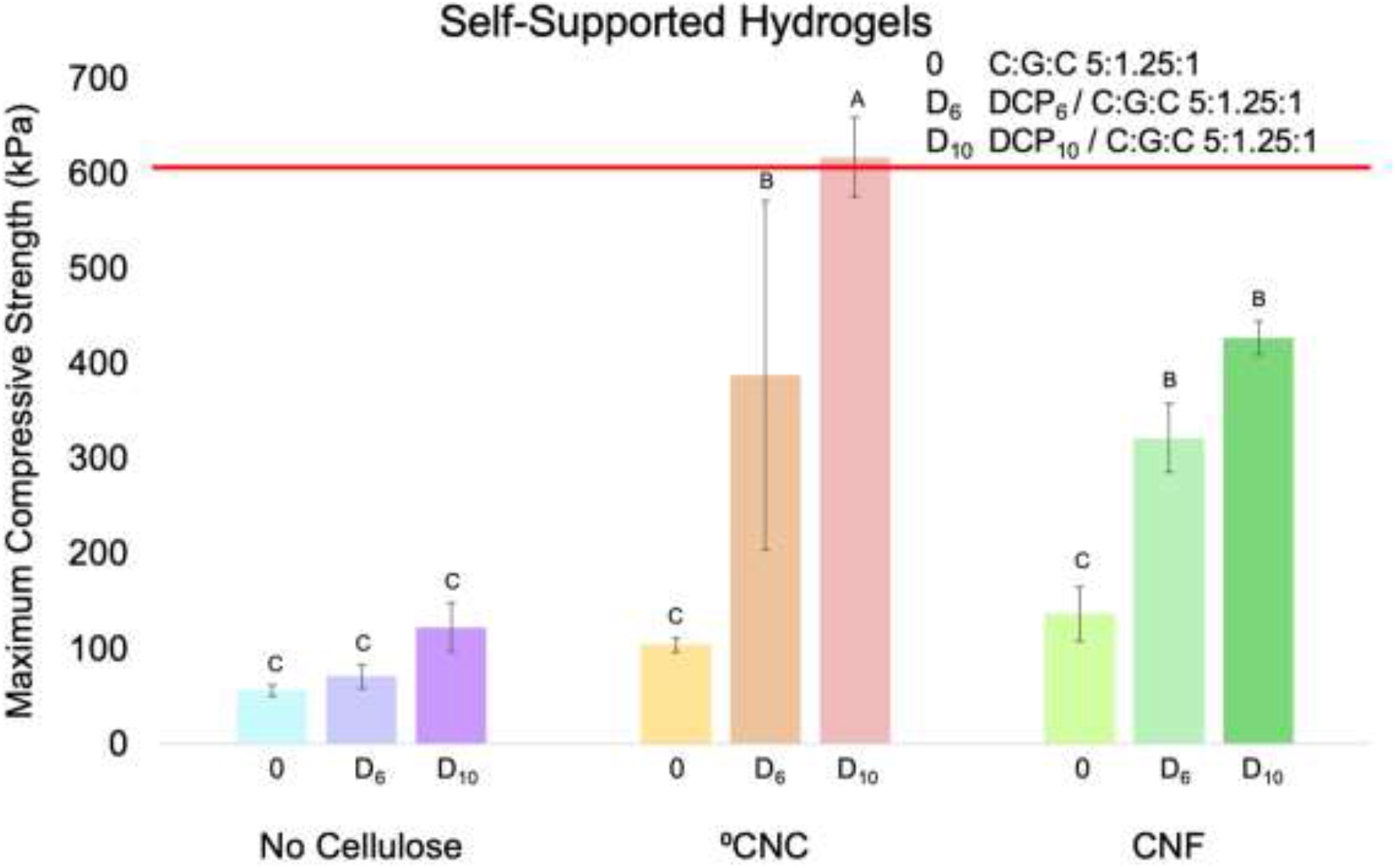
Compression strength for C:G:C 5:1.25:1 hydrogels formed with various cellulose types and DCP concentrations. Hydrogels were prepared with combinations of no cellulose, ^0^CNCs, or CNFs, without DCP (0), with 6% DCP (D_6_), or with 10% DCP (D_10_). The lower bound of the compressive strength of vertebral bone is indicated with a red line at 600 kPa. Values are reported as the average ± standard deviation (N = 4). Statistical groupings are based on a Tukey’s HSD test performed using all groups shown. Groups that possess different letters have statistically significant differences (p < 0.05) in their means whereas those that possess the same letter are statistically similar.

While CNF incorporation improved the compressive strength of C:G:C hydrogels, these formulations still fall below the desired minimum threshold of 600 kPa. A potential way to improve the mechanical strength of hydrogels is to harness the structural integrity of bone itself. To achieve this, chitosan hydrogels were formed around and within crushed cancellous chip allograft (AG). Interestingly, in contrast to the small amount of water exclusion observed during self-supported hydrogel formation, allograft-embedded hydrogels retain the full solution volume used for their synthesis. When tested for their compressive strength, allograft incorporation resulted in a considerable increase in compressive strength (**Figure 6**) when compared to just the inclusion of cellulose and/or DCP into the hydrogel (**Figure 5**). This influence overwhelmed the synergistic impact of cellulose type and/or DCP incorporation into allograft-embedded hydrogels except for the case of the CNF / DCP_6_ / C:G:C formulation. In addition to having the highest compressive strength, the CNF / DCP_6_ / C:G:C hydrogels tested had remarkedly similar values yielding a very low variance resulting in statistical significance when compared to the CNF / C:G:C hydrogels. Due to the much higher standard deviations calculated with all other allograft-embedded hydrogels, it is much more likely that the CNF / DCP_6_ / C:G:C hydrogel samples evaluated did not accurately represent the entire possible sample population rather than a true innate difference existing with this formulation. The improvement in compressive strength measured for allograft-embedded hydrogels provides substantial support for their use as a biomaterial for bone regeneration applications though their biocompatibility and osteoinductivity still had to be probed *in vitro* before these could ever be assessed *in vivo*.

**Figure 6.**
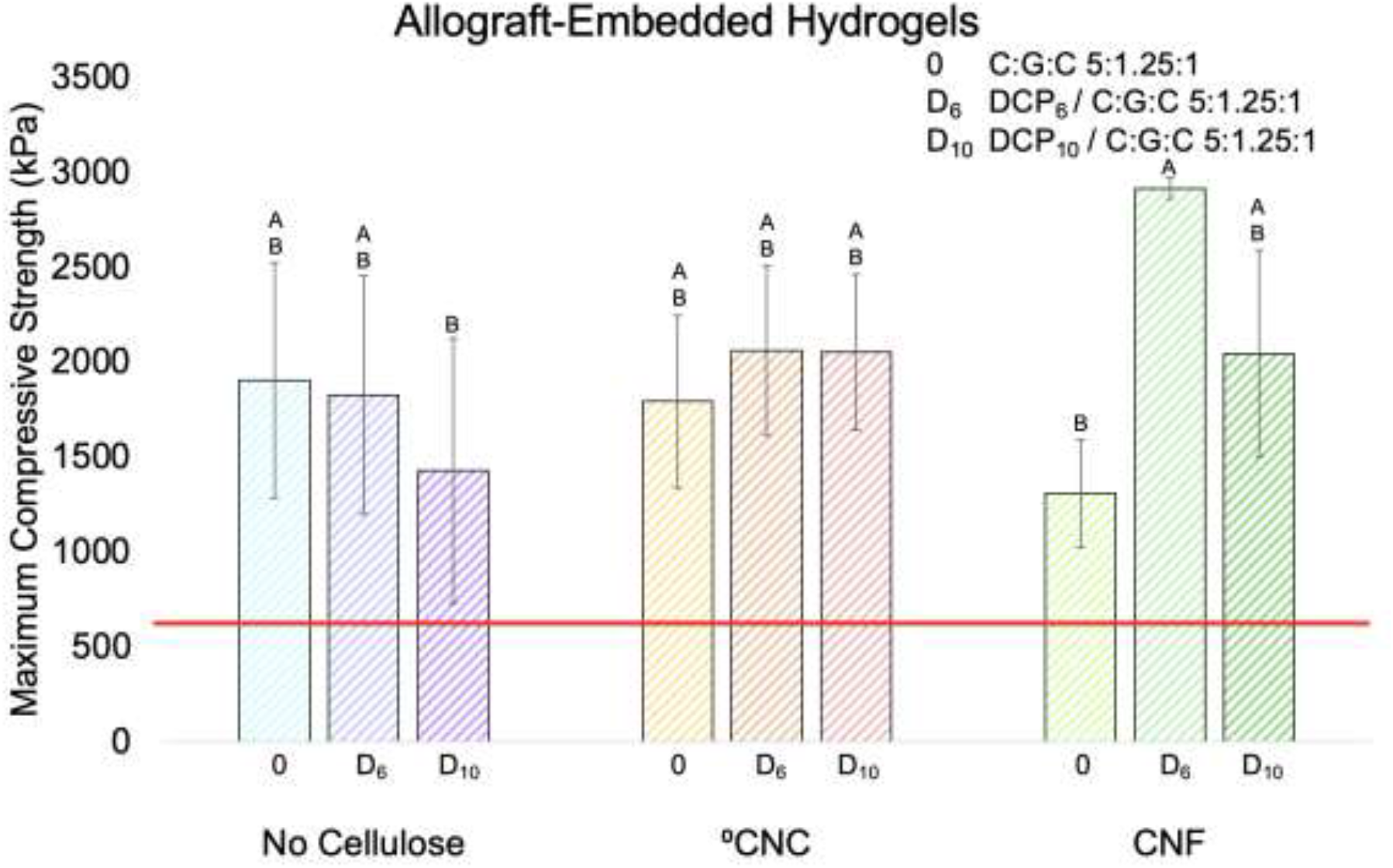
Compression strength for allograft-embedded C:G:C 5:1.25:1 hydrogels with various cellulose types and DCP concentrations. Hydrogels were prepared with combinations of no cellulose, ^0^CNCs, or CNFs, without DCP (0), with 6% DCP (D_6_), or with 10% DCP (D_10_) all gelated around crushed cancellous chip allograft. The lower bound of the compressive strength of vertebral bone is indicated with a red line at 600 kPa. Values are reported as the average ± standard deviation (N = 4). Statistical groupings are based on a Tukey’s HSD test performed using all groups shown. Groups that possess different letters have statistically significant differences (p < 0.05) in their means whereas those that possess the same letter are statistically similar.

### Impact of Agent Incorporation on Chitosan Hydrogels Biological Properties

With the effects DCP content, cellulose type, and/or allograft inclusion have on chitosan hydrogel material properties established, their biocompatibility and bioactivity were able to be studied *in vitro* using murine mesenchymal stem cells (mMSCs). To further investigate the potential negative biological side effects observed previously,^8,9^ an initial 24-hour cytotoxicity study was performed employing extract media from self-supporting hydrogels prepared with combinations of no cellulose, ^0^CNCs, or CNFs, without DCP (0), with 6% DCP (D_6_), or with 10% DCP (D_10_). The cytotoxic impact of these biomaterials was determined by comparing the quantity and total metabolic activity of hydrogel extract treated mMSCs using the Quant-iT™ PicoGreen™ dsDNA assay (**Figure 7A**) and an alamarBlue™ assay (**Figure 7B**), respectively, to cells grown in complete growth media alone. Within the no cellulose and CNF hydrogel groups, DCP incorporation had a negative impact on the cells. As this media was transferred to the cells, they would have been exposed to a bolus of the full 24-hour dose of hydrogel dissociated components which could have easily overwhelmed the cells. Interestingly, the ^0^CNC hydrogels were found to be so cytotoxic that the impact of DCP incorporation could not be effectively studied with these materials. The negative biological impact observed with ^0^CNC-containing hydrogels is likely attributable to the long alkyl chain associated with the previously mentioned CTAB charge capping group used to generate ^0^CNCs. This is supported by previous research that found un-bound or detached CTAB can be a potent source of cytotoxicity.^20,21^ The incorporation of CNFs into hydrogels facilitated promising biocompatibility which is unsurprising given the excellent biocompatibility that is inherent to unmodified cellulose.^22,23^ This result further supports CNFs as a promising alternative to ^0^CNCs. Therefore, ^0^CNC hydrogels were excluded from further biological testing. Hydrogels with no cellulose as well as the CNF / DCP_6_ / C:G:C formulation were also not further tested. While hydrogels with no cellulose possessed promising biocompatibility, they did not exhibit the mechanical properties necessary to be used for vertebral bone regeneration applications (**Figure 5**). Additionally, since CNF / DCP_6_ / C:G:C and CNF / DCP_10_ / C:G:C hydrogels had indistinguishable indicators of osteoinductive potential (*i.e*., SSM release profile and acute biocompatibility), CNF / DCP_10_ / C:G:C hydrogels were chosen because they displayed the potential to be a more mechanically competent formulation (**Figure 5**). Hydrogels with and without AG were also studied due to the considerable enhancement of compressive strength they were able to achieve (**Figure 6**) and the fact that AG is commonly used in spinal fusion procedures.^24,25^

**Figure 7.**
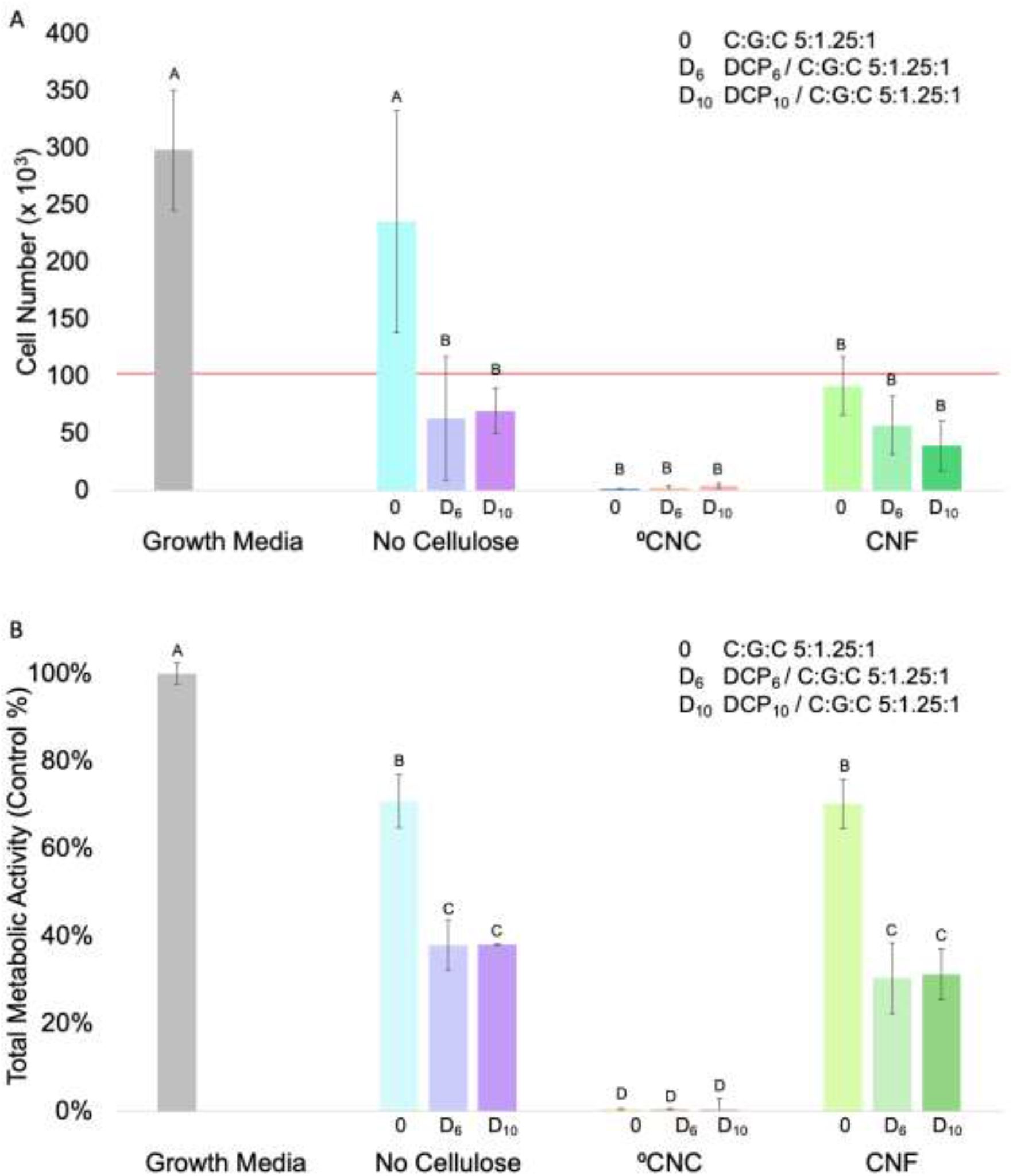
Biocompatibility of self-supported C:G:C 5:1.25:1 hydrogels formed with various cellulose types and DCP concentrations. Hydrogels were prepared with combinations of no cellulose, ^0^CNCs, or CNFs, without DCP (0), with 6% DCP (D_6_), or with 10% DCP (D_10_). (**A**) Cell proliferation and (**B**) total metabolic activity were measured for cells cultured for 24 hours in extract media at 37°C. As 24-well plates were seeded with 100,000 cells, this is indicated by a red line. Cell number was determined via a Quant-iT™ PicoGreen™ dsDNA assay. Total metabolic activity was assessed via an alamarBlue™ assay and standardized to control cells cultured in growth media. Data is reported as the mean ± standard deviation (N = 4). Statistical groupings are based on a Tukey’s HSD test performed using all groups shown. Groups that possess different letters have statistically significant differences (p < 0.05) in means whereas those that possess the same letter are statistically similar.

To evaluate the long-term biocompatibility of the CNF / C:G:C hydrogels, mMSCs were exposed to hydrogels prepared with and without 10% DCP (DCP_10_) formed with and without AG. Complete growth media and osteogenic media as well as DCP_10_ and allograft delivered via a semi-permeable insert were each used as controls for the hydrogels to evaluate their individual effects on the cells. Biocompatibility was evaluated by measuring the cell proliferation (**Figure 8A**) and total metabolic activity (**Figure 8B**) at 3, 7, 14, and 21 days. DCP_10_ or AG present in semi-permeable inserts had little effect on the quantity and viability of exposed cells at any timepoint compared to growth media. In contrast, cells grown in osteoinductive media lagged in their cell number and viability, suggesting they may have undergone differentiation into osteoblasts.^26^ For all hydrogels investigated, exposed cell quantity and viability were significantly lower than all control groups at day 3, indicating some initial toxicity and/or proliferative suppression below the 50,000 starting cell number (*i.e*., averages of 12,840 - CNF / C:G:C, 10,870 - CNF / DCP_10_ / C:G:C, 43,560 - AG / CNF / C:G:C, and 26,500 - AG / CNF / DCP_10_ / C:G:C). This data parallels the 24-hour cytotoxicity results presented in **Figure 7**. Interestingly, hydrogels with AG (*i.e*., AG / CNF / C:G:C and AG / CNF / DCP_10_ / C:G:C) less negatively impacted cell numbers than those without AG (*i.e*., CNF / C:G:C and CNF / DCP_10_ / C:G:C) though incorporating DCP within allograft-embedded hydrogels led to similar suppression seen in **Figure 7** and made this group have a statistically insignificant difference in its cell count when compared to the self-supported hydrogels. This behavior could be due to highly porous AG acting as a sink and preventing the rapid expulsion of hydrogel components that are believed to have played a role in overwhelming the cells at 24 hours (**Figure 7**). At days 7, 14, and 21, cells exposed to the two hydrogels with AG followed a similar growth pattern as the osteogenic media lagging behind the other three controls (*i.e*., growth media, DCP_10_, and AG). Even though the CNF / C:G:C and CNF / DCP_10_ / C:G:C hydrogels suppressed cell growth and viability at day 3, cells exposed to these formulations recovered their quantity and total metabolic activity to those cultured in osteoinductive media by days 7 and 14, respectively. A reason for the later-stage growth is that the hydrogels dissociate rapidly in the first 3 days and stabilizes thereafter (**Figure 3**). Therefore, the cells were likely exposed to a bolus of hydrogel components initially which then slowed after the 3 day window allowing cell populations to recover and grow. The retention of the proliferative capacity of healthy mMSCs when exposed to CNF / C:G:C hydrogels is a promising improvement over our previous °CNC / chitosan hydrogel formulations.^8,9^

**Figure 8.**
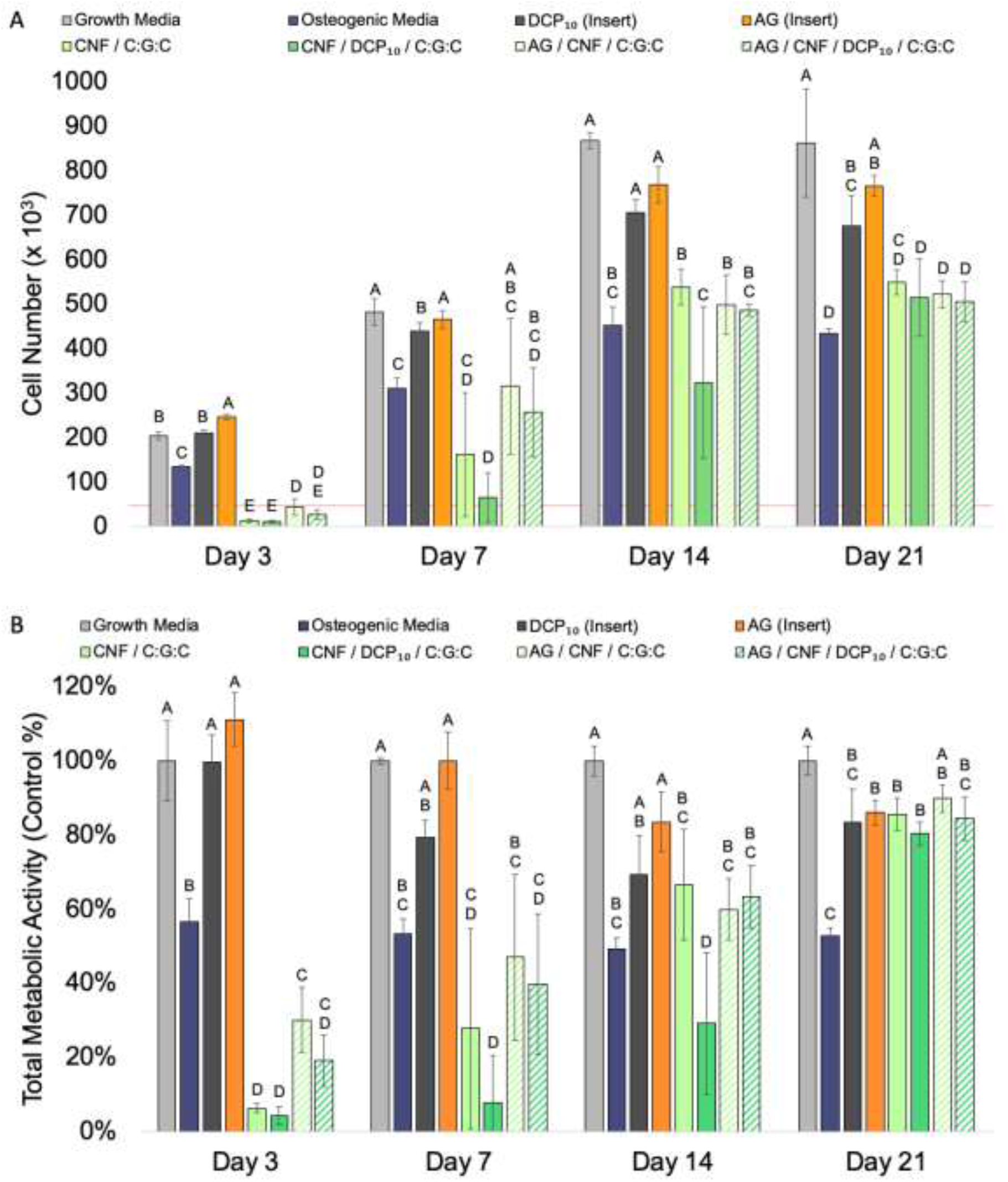
Biocompatibility of self-supported and allograft-embedded CNF / C:G:C hydrogels formed with various DCP concentrations. Hydrogels were prepared with or without DCP_10_ and with or without allograft (AG). (**A**) Cell proliferation and (**B**) metabolic activity were measured for cells cultured for up to 21 days in growth media at 37°C. Cell number was determined via the Quant-iT™ PicoGreen™ dsDNA assay. Total metabolic activity was assessed via an alamarBlue™ assay and standardized to control cells cultured in growth media. Values are reported as the average ± standard deviation (N = 4). Statistical groupings are based on a Tukey HSD comparison between groups at the same timepoint. Groups that possess different letters have statistically significant differences (p < 0.05) in means whereas those that possess the same letter are statistically similar.

To evaluate the osteoinductivity of CNF / C:G:C hydrogels, alkaline phosphatase (ALP) and alizarin red (ALZ) assays were performed on mMSCs cultured with formulations prepared with or without 10% DCP (DCP_10_) and with or without AG over 21 days (**Figure 9**). Exposing cells to DCP_10_ or AG delivered in semi-permeable inserts had minimal to mild effects on their ALP production whereas those cultured in osteogenic media had statistically significantly higher ALP synthesis at each time point when compared to mMSCs grown in growth media (**Figure 9A**). While a lack of ALP activity when exposed to DCP_10_ seems to contradict our previous results,^8-10^ the Ca^2+^ and P_i_ release concentrations observed in this most recent work differs significantly from our earlier efforts. Specifically, the DCP tested here released high Ca^2+^ and low P_i_ concentrations near the edges of the therapeutic window likely limiting its inductive capacity. A possible explanation for this discrepancy is the DCP used for this work was from a different batch than the one employed previously though further studies would need to be conducted to explore this effect further. All hydrogel formulations tested induced cells to produce high levels of ALP similar or greater to osteogenic media at day 3. The higher numbers observed with self-supported hydrogels could be related to the very low cell numbers found within those wells (**Figure 8A**). Interestingly, hydrogel-exposed mMSCs saw their ALP production return to baseline levels at the later time points in the study (*i.e*., days 7, 14, and 21). As ALP is an early marker of osteoinduction, the results observed for the hydrogel groups may suggest their co-cultured mMSCs are differentiating down an osteogenic lineage.^27^

**Figure 9.**
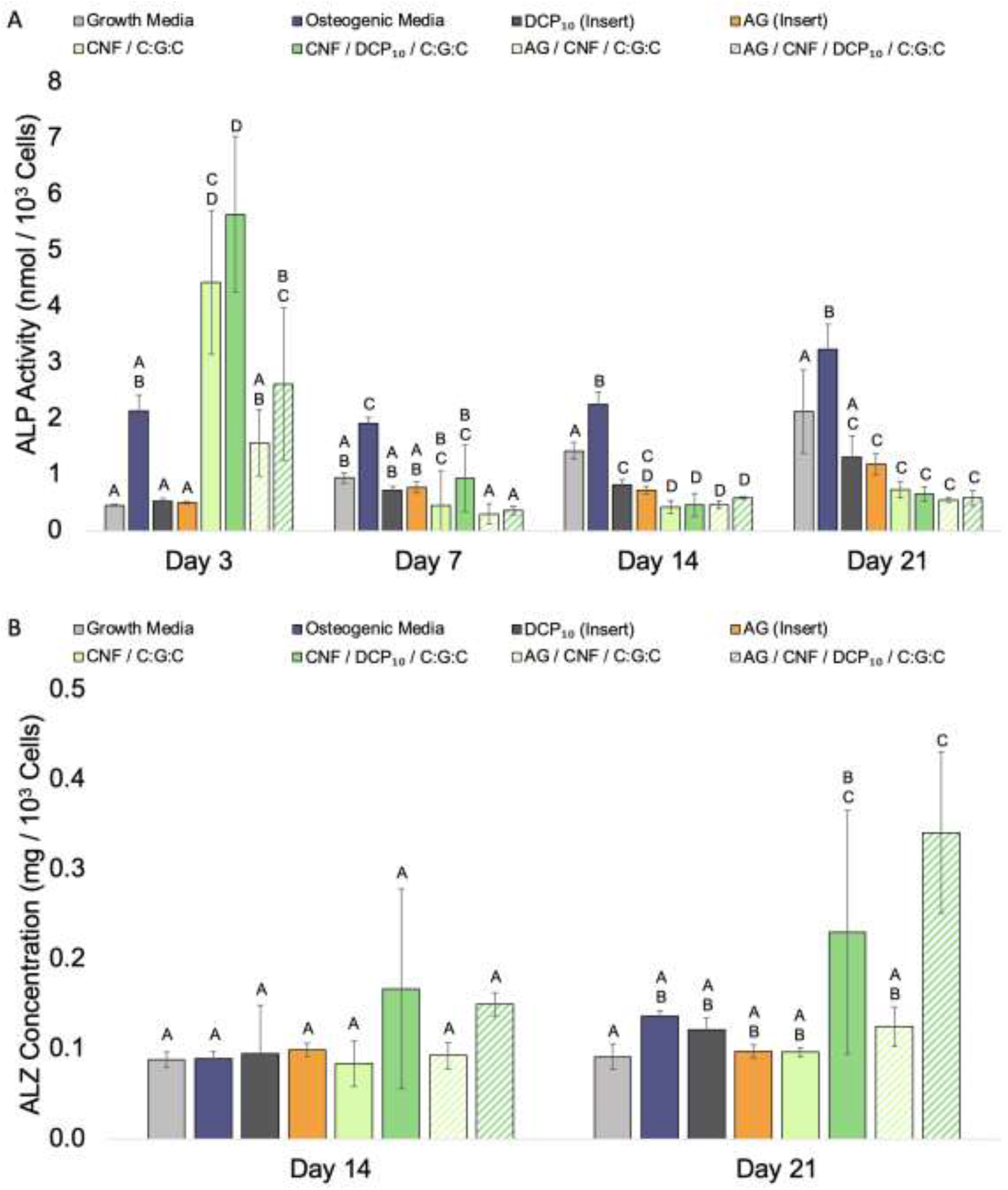
Osteoinductivity of self-supported and allograft-embedded CNF / C:G:C hydrogels formed with various DCP concentrations. Hydrogels were prepared with and without DCP_10_ and with or without allograft (AG). (**A**) ALP activity and (**B**) cell-based mineralization were measured for cells cultured for up to 21 days in growth media at 37°C. ALP activity was determined via an ALP pNPP assay. Alizarin red (ALZ) staining was used as an indirect measure of mineralization with and without hydrogels. ALZ content for matching acellular hydrogel formulations over the same incubation time was subtracted to determine cell-base mineralization. Data is reported as the mean ± standard deviation (N = 4). Statistical groupings are based on a Tukey’s HSD test performed using all groups at the same timepoint. Groups that possess different letters have statistically significant differences (p < 0.05) in means whereas those that possess the same letter are statistically similar.

To complement this early-stage osteoinductivity data, late-stage mineralization as evidenced by CaP ceramic deposition, was studied. This was assessed at the latter two timepoints (*i.e*., days 14 and 21) via extracellular matrix fixed calcium staining using an Alizarin Red (ALZ) assay (**Figure 9B**). For the groups that showed elevated ALP activity (*i.e*., osteogenic media and DCP-containing hydrogels), interesting differences in their ALZ results were observed. Osteogenic media did not elevate cell-based mineralization above growth media at either day 14 or day 21. As ALP activity remained elevated through these time points, it is likely that exposure to the osteogenic media was able to initiate osteogenic differentiation, but not facilitate the mMSCs to become full-fledged osteoblasts, a result consistent with previously published research.^28,29^ In contrast, though cells exposed to the two DCP-containing hydrogels had background levels of mineralization at day 14, they both had modestly elevated ALZ content significantly greater than mMSCs cultured in growth media. While the overall magnitude of ALZ content on a per cell basis was lower than previously reported,^8,9^ the much greater cell numbers seen in this more recent research compensated for this deficiency. Specifically, the analogous hydrogel (*i.e*., °CNC / DCP_10_ / C:G:C 5:2.5:1) from our published work^9^ induced ∼ 151 mg of ALZ-stained mineral compared to ∼ 119 mg and ∼ 172 mg caused by exposure to CNF / DCP_10_ / C:G:C 5:2.5:1 hydrogel and AG / CNF / DCP_10_ / C:G:C 5:2.5:1 hydrogel, respectively. The combination of early ALP expression and mineralization from cells exposed to DCP-containing hydrogels indicates that these biomaterials are indeed osteoinductive.

### Evaluation of Chitosan Hydrogels *in Vivo* Biological Properties

To build upon the exciting *in vitro* bioactivity observed with composite hydrogels, their *in vivo* biocompatibility and spinal fusion properties were assessed. ^15,30-33^ First, various hydrogel formulations were implanted intramuscularly into C57BL/6 mice for 14 days. All mice were found to be healthy through the 14 days of the study. The animals remained bright, alert, and responsive while possessing normal eating and drinking habits as well as retaining full function of the limb operated upon. Upon examination of the implant site after euthanasia, all surrounding tissues were evaluated for signs of infection and inflammation in which no differences were found among the test groups. Harvested tissue was fixed, sectioned, and stained with hematoxylin and eosin (H&E) stain and evaluated for various cell types by a pathologist blinded to the formulation used and physiological responses were recorded in accordance with ISO 10993 – 6. Examination criteria and the results of this study are provided in **Table 2** and **Table 4**, respectively.

**Table 4.** Histology scoring for implanted CNF / C:G:C hydrogels after 14 days. Sterilized (*) and unprocessed CNF / C:G:C hydrogels were prepared with and without 10% DCP (DCP_10_) and with or without crushed cancellous chip allograft (AG) (N = 4). Values reported as the average score of each group.

|  | PMN | Lymphocytes | Plasma | Macrophages | Giant | Necrosis | Neovascularization | Fibrosis | Fatty Infiltrate | Traumatic necrosis | Foreign debris | Total |
| --- | --- | --- | --- | --- | --- | --- | --- | --- | --- | --- | --- | --- |
| CNF / C:G:C | 2 | 1 | 0 | 2 | 0 | 0 | 2 | 0 | 0 | 0 | 0 | 7 |
| CNF / C:G:C* | 2.6 | 1.2 | 0 | 1.2 | 0 | 0 | 1.4 | 0 | 0 | 0.6 | 0 | 7 |
| CNF / DCP<sub>10</sub> / C:G:C | 2.4 | 1.4 | 0 | 1.8 | 0 | 0 | 1.4 | 0 | 0 | 0 | 0 | 7 |
| CNF / DCP<sub>10</sub> / C:G:C* | 3 | 1.6 | 0 | 2.2 | 0.4 | 0 | 2.2 | 0 | 0 | 0 | 0 | 9.4 |
| AG / CNF / C:G:C | 3 | 1.2 | 0 | 1 | 0 | 0 | 2 | 0 | 0 | 0 | 0 | 7.2 |
| AG / CNF / C:G:C* | 2.25 | 2 | 0 | 1 | 0 | 0.25 | 1.75 | 0 | 0 | 0 | 0 | 7.25 |
| AG / CNF / DCP<sub>10</sub> / C:G:C | 2.8 | 1.2 | 0 | 1.2 | 0 | 0.6 | 2.2 | 0 | 0 | 0 | 0 | 8 |
| AG / CNF / DCP<sub>10</sub> / C:G:C* | 2.6 | 1.6 | 0 | 1.6 | 0 | 0 | 2 | 0 | 0 | 0 | 0 | 7.8 |

Detailed conclusions are difficult to draw from the histological assessment data given that the control was nearly inert to the surrounding tissue and was unable to be evaluated for scoring due to the lack of an identifiable physical implant. However, we can still learn about the ongoing physiological processes at day 14 based on the cell populations present in the surrounding tissues. Across all groups, there was a greater presence of polymorphonuclear cells (i.e., PMNs) such as neutrophils which are an early indicator of the foreign body response. There was a slightly lower expression of chronic inflammation indicators such as lymphocyte and macrophage infiltration as well as no local presence of plasma cells. This suggests that the tissue is still in the acute phase of the foreign body response with only modest indications of chronic inflammation. The 14-day biocompatibility study revealed that the complex chitosan hydrogels evaluated were non-toxic and therefore cleared for study in higher order animal in vivo studies like the rabbit spinal fusion model.

To evaluate the application of these hydrogels for osteomodulatory applications, CNF / DCP_10_ / C:G:C and AG / CNF / DCP_10_ / C:G:C hydrogels were compared to the implantation of AG alone in a model on New Zealand White Rabbits. Assessment scores from 2D radiographs taken 0, 3, and 6 weeks post-operation are presented in **Figure 10**. All three groups were found to be radiopaque immediately after surgery as indicated by their non-zero score at this timepoint. Additionally, all three groups facilitated enhanced neotissue mineralization over the 6 weeks of the study though differences were observed by the end of the experiment. CNF / DCP_10_ / C:G:C hydrogels induced the highest fusion score (3.67 ± 0.30), followed by AG / CNF / DCP_10_ / C:G:C hydrogels (3.06 ± 0.90), and AG alone was rated the lowest (2.44 ± 1.11). While statistical analysis was not performed due to the semi-quantitative nature of this assessment, the data still provides evidence that the CNF / DCP_10_ / C:G:C hydrogels prompted the most rapid new bone formation as determined by 2D radiography. This finding is supported by the osteoinductive nature of this hydrogel determined *in vitro* especially in contrast to AG alone (**Figures 8 & 9**). These results agree with the literature which reports that allografts are generally found to be non-osteoinductive and result in lower fusion rates when used alone.^6^ For this reason, they are commonly compounded with proteinaceous growth factors or used in conjunction with autografts.^6,34^ When we combined allograft with our osteoinductive system to yield AG / CNF / DCP_10_ / C:G:C hydrogel, a slight improvement in new bone formation over the AG alone was observed. Though promising, this induced less neotissue development than the CNF / DCP_10_ / C:G:C hydrogel suggesting the ability of our novel product to expedite spinal fusion outcomes without the incorporation of allogeneic materials.

**Figure 10.**
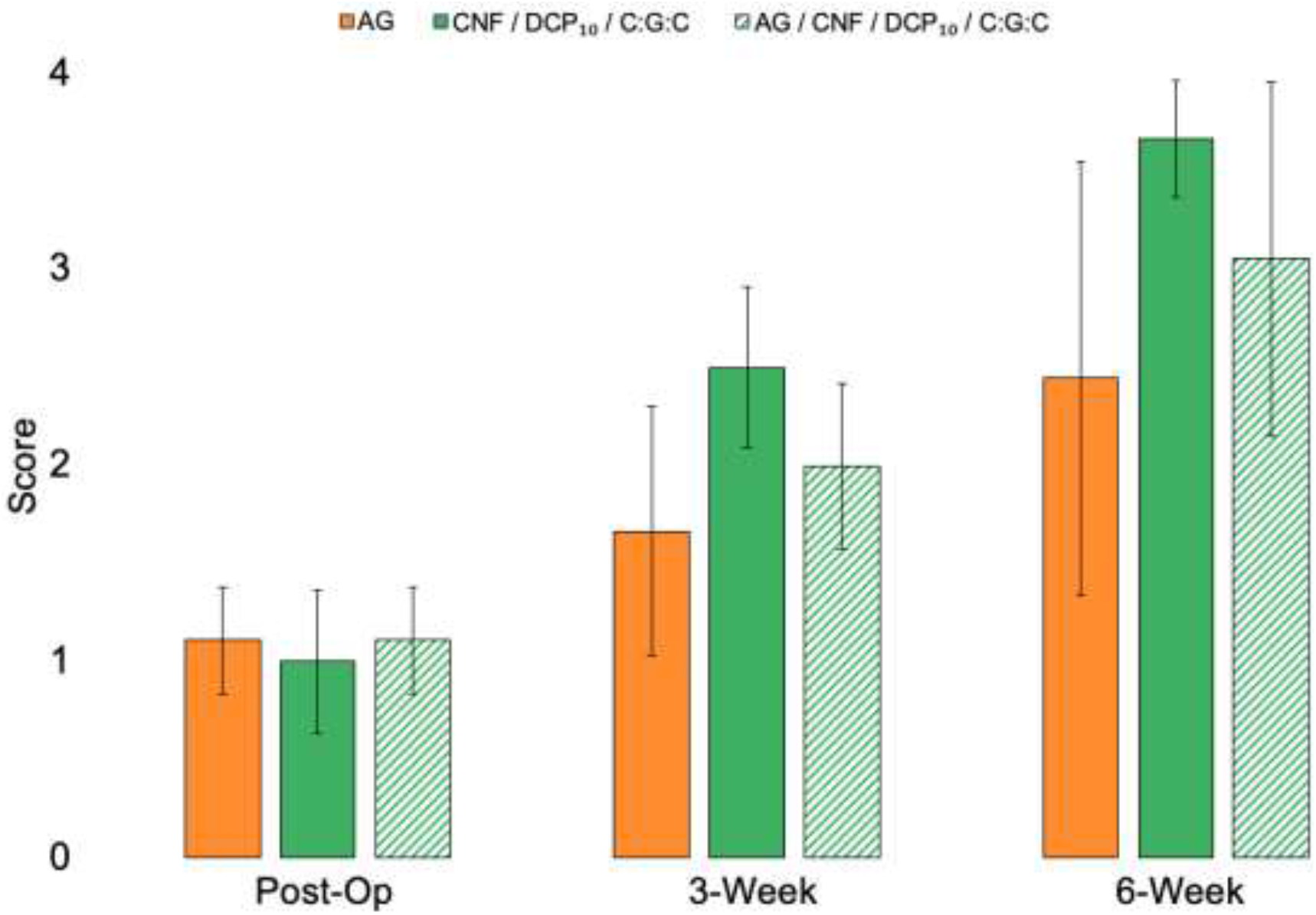
2D radiograph scoring of implanted spinal fusion materials of images taken immediately post operation, at 3-weeks and 6 weeks post-operation. Three graft materials (*i.e*., AG, AG / CNF / DCP_10_ / C:G:C, CNF / DCP_10_ / C:G:C) were evaluated for their ability to achieve fusion in a posterolateral intertransverse process lumbar fusion performed on New Zealand White Rabbits. Preliminary data suggests that all groups were able to induce some amount of new mineralized tissue between the vertebral levels.

## Conclusions

The materials and biological properties determined for the CNF / DCP_10_ / C:G:C and AG / CNF / DCP_10_ / C:G:C hydrogels show tremendous promise for bone tissue engineering applications. Specifically, these formulations combine desirable gelation time, swellability, resistance to dissociation, ion release, and mechanical strength with biocompatibility and osteoinductivity, outcomes that were only achievable by incorporating CNFs, DCP, and AG into chitosan hydrogels. While work previously conducted by our group explored chitosan hydrogels for the controlled release of osteoinductive ions for bone regeneration, significant mMSC proliferation suppression was observed with these formulations. Before these biomaterials can be translated for clinical applications, they needed to be further modified and optimized to alleviate this issue. This research has shown that by substituting the mechanically reinforcing agent from cytotoxic ^0^CNCs to more biocompatible CNFs, hydrogels could be fabricated that possess similar desirable mechanical properties, but with improved biological properties. Additionally, AG incorporation into the hydrogels resulted in even greater mechanical strength, biocompatibility, and osteoinductivity when compared to the allograft-free hydrogels. When evaluated *in vivo* for biocompatibility, the hydrogels were found to be non-toxic though the subacute phase (14 days). A 6-week spinal fusion study deploying the hydrogels found them to achieve near complete fusion over this time period when evaluated radiographically. These results suggest that multi-component loaded chitosan hydrogels can not only serve as an osteoinductive biomaterial for spinal fusion applications, but can be potentially expanded upon to treat vertebral compression fractures, traumatic bone fractures, and segmental bone defects.

## Acknowledgments

This research was conducted with help from numerous individuals that should be recognized. Dr. Sarah Schlink, DVM and Dr. Samantha Gerb, DVM were instrumental in ensuring all *in vivo* models were conducted in compliance with University of Missouri Animal Care and Use Committee standards. Ashley Szczordroski provided considerable assistance in conducting the MicroCT scans of the spinal tissue. A special thanks goes to Dr. Grant Van Dyke Darkow, MD for his evaluation and scoring of the murine biocompatibility histological slides. Dr. Wellington Hsu, MD and his team at Northwestern University were kind enough to train our team in the rabbit spinal fusion model. This work was supported and funded through a MTF Biologics Junior Investigator Extramural Research Grant and a MU Coulter Bridge Grant.

